# CNHplus: the chromosomal copy number heterogeneity which respects biological constraints

**DOI:** 10.1101/2022.09.30.510279

**Authors:** Marian Grendár, Petr Martínek, Dušan Loderer, Ondrej Ondič

## Abstract

**Motivation:** Intra-Tumor Heterogeneity (ITH) is a hallmark of cancer. ITH influences evolution of tumors and hence also affects patients survival. There are several methods for quantifying ITH. Recently, van Dijk et al. (*Nat. Commun*. 12(1):1-12, 2021) introduced Copy Number Heterogeneity (CNH) method for quantifying ITH from a single tumor sample, based on a copy number profile. The authors demonstrated capability of CNH to stratify patients for survival, across cancers. We aimed to reproduce the authors’ study.

**Results:** A deficiency in CNH is pointed out. The absolute copy number (ACN) profile obtained by solving the CNH optimization problem may contain negative number of copies. For a large portion of samples from The Cancer Genome Atlas (TCGA) studies considered by the authors the CNH-recovered ACN profiles are faulty. CNHplus corrects the flaw by imposing the non-negativity constraint. CNHplus is applied to survival stratification of patients from the TCGA studies. Also, it is discussed which other biological constraints should be incorporated into CNHplus. CNHplus estimates of the tumor purity, tumor ploidy are compared with those of the ABSOLUTE method.

**Availability and implementation:** R library CNHplus can be accessed here: https://github.com/grendar/CNHplus

## 1 INTRODUCTION

Cancer cells are not exempt from evolutionary pressure. Intra-Tumor Heterogeneity (ITH) (cf. McGranahan and Swanton (2015), Li, Seehawer, and Polyak (2022)) of a cancer cell population reflects adaptation of cancer cells to new circumstances; cf. Van Den Bosch et al. (2021). For this reasons a biomarker of ITH should in principle be able to stratify patients for survival, as patients with the high ITH should be expected to have poorer survival than patients with the low ITH.

Several methods for quantifying ITH were developed over the recent years. Van Den Bosch, Vermeulen, and Miedema (2022) propose a categorization of the methods and provide a survey of the mathematical models that are behind the methods. There are single-sample methods as well as multi-sample methods, with their advantages and disadvantages. Many of the methods are based on variant allele frequencies. More recent are methods that utilize copy number variations. One of them is the Copy Number Heterogeneity (CNH) method developed by van Dijk et al. (2021). CNH uses a single tumor sample to quantify the extent to which an Absolute Copy Number (ACN) profile of a tumor differs from the values 0,1,2, … that correspond to homogeneous populations. The greater the CNH, the greater should be the genetic heterogeneity in the cancer cell population; cf. Van Den Bosch et al. (2021). And with the greater heterogeneity of the malignant cell population should be associated the poorer survival of a patient. van Dijk et al. (2021) use The Cancer Genome Atlas (TCGA) studies (cf. Weinstein et al. (2013), Liu et al. (2018)) to demonstrate that CNH is able to stratify patients for survival in many cancers.

For a Relative Copy Number (RCN) profile, van Dijk et al. (2021) obtain CNH by searching over a grid of pairs of values of tumor purity and tumor ploidy for such a pair, for which the associated ACN of tumor is closest to the integer values. The recovered ACN profile of tumor should, in our view, satisfy biological constraints. We point out that restricting tumor purity and tumor ploidy in the CNH computation to biologically meaningful values does not, in general, guarantee that the recovered ACN profile of tumor is nonnegative in every segment. There may be segments with a negative number of copies. For instance, in TCGA-UCEC study, out of the 540 samples from the study that were considered by van Dijk et al. (2021), 434 are such, that the recovered ACN profile of the tumor has at least one segment with the negative number of copies. We introduce CNH+, the corrected CNH method, which respects the non-negativity of the ACN profile of tumor and analyze TCGA data.

## 2 DEFINITION OF CNH

In order to make this note selfstanding, the definition of CNH is repeated here, using the terminology, notation and equations numbering of van Dijk et al. (2021), at the price of obtaining disordered numbering of equations. Equations below, with non-numeric labels, are introduced here to provide necessary notions.

Bioinformatic copy number calling usually results in the RCN profile *r* of a sample, where the relative copy number *r_i_*, of segment *i* with width *w_i_*, can be seen as the ratio of the ACN of the sample in the segment to the average ACN of the sample across the entire genome. Interest however lays in the ACN profile of the tumor, rather than the sample. Assuming the single dominant clone in the tumor, the sample can be seen as a mixture of a proportion *α* of cancer cells and 1 – *α* of normal diploid cells. Then the RCN profile of the sample *r* and the ACN profile *q* of the tumor are related as (cf. Carter et al. (2012)):

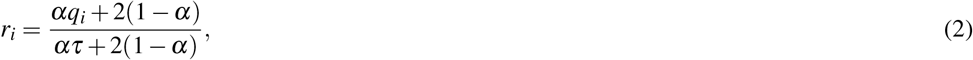

where *α* is the tumor purity and *τ* is the average tumor ploidy, which is defined as

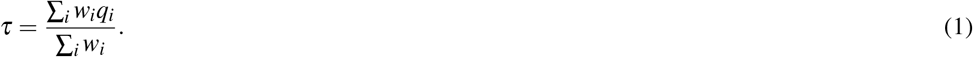

From the RCN profile *r* of the sample it is desired to estimate the tumor purity *α*, tumor ploidy *τ* and the associated ACN profile *q* of the tumor. There are several methods for estimating these quantities of biological interest; to a large extent they arise from the seminal work Carter et al. (2012).

van Dijk et al. (2021) estimate the tumor purity *α* and the average tumor ploidy *τ* as points of minima of the function *κ*(*α, τ, r, w*) which is defined as

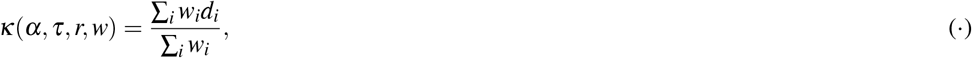

where the *i*-th component *d_i_*, of *d* is defined as

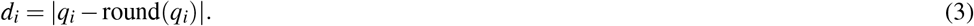

Formally put, the van Dijk et al. (2021) estimates 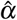 of *α* and 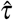 of *τ* are given as

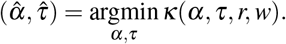

Note that *q* can be expressed from (2) as

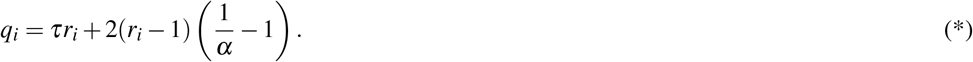

Plugging 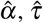 and *r* into (*) leads the estimate 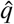 of the ACN profile *q* of tumor.

CNH of the tumor is 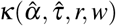, i.e., the value of the function *κ*(*α, τ, r, w*) at its minimum over (*α, τ*). Or equivalently in words of van Dijk et al. (2021),

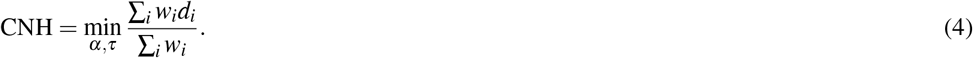

CNH is the minimum of the weighted *L*_1_ distance between *q* and round(*q*), where the minimization is carried over biologically relevant ranges of *α* and *τ*. The authors suggest *α* ∈ [0.2,1] and *τ* ∈ [1.5,5].

### 2.1 Invariance of CNH

The value of *κ* is invariant to a shift of *q*. Adding an integer to ACN profile *q* does not change the value of CNH. If *q* is shifted by *c* up or down to *q*′, then it follows from (1) that the average tumor ploidy *τ* changes to *τ*′ = *τ* + *c*. From (*) then follows that the tumor purity *α* changes to 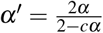. Hence, (*α, τ, q*) and (*α*′, *τ*′, *q*′) have the same value of CNH. Since *α*′, *τ*′ can take values from biologically relevant ranges, only, not every shift of *q* is possible.

Stated from a different angle - there are samples for which the solution pair 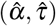 need not be unique. In such a case, the candidate solutions lead different ACN profiles with the same value of CNH. And a method for disambiguation of the candidate solutions is needed, in order to obtain the CNH estimates of the tumor purity, the tumor ploidy and ACN profile of tumor. Sauer et al. (2021), who solve the same problem (4) as van Dijk et al. (2021) suggest using additional information (such as mutant allele fraction of TP53) to disambiguate the competing solutions.

## 3 ABSOLUTE COPY NUMBER PROFILE OBTAINED BY CNH OPTIMIZATION

Estimated ACN profile 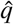 of the tumor, which is associated with the estimated tumor purity 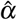 and estimated tumor ploidy 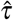, should satisfy biological constraints. In particular, the number of copies 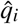 of segment *i* should be non-negative, for every *i*.

Unfortunately, restricting (*α, τ*) in the CNH optimization to biologically meaningful values does not, in general, guarantee that the recovered ACN profile will be non-negative in every segment. The next section illustrates the flaw in CNH.

## 4 EXAMPLE OF THE FLAW IN CNH

Consider for instance the TCGA Uterine Corpus Endometrial Carcinoma (UCEC) study; cf. Berger et al. (2018). The RCN profile for one of the samples from the study (TCGA-AJ-A3EJ-01A-11D-A19X-01) is plotted on Fig. 1.

**Figure 1.**
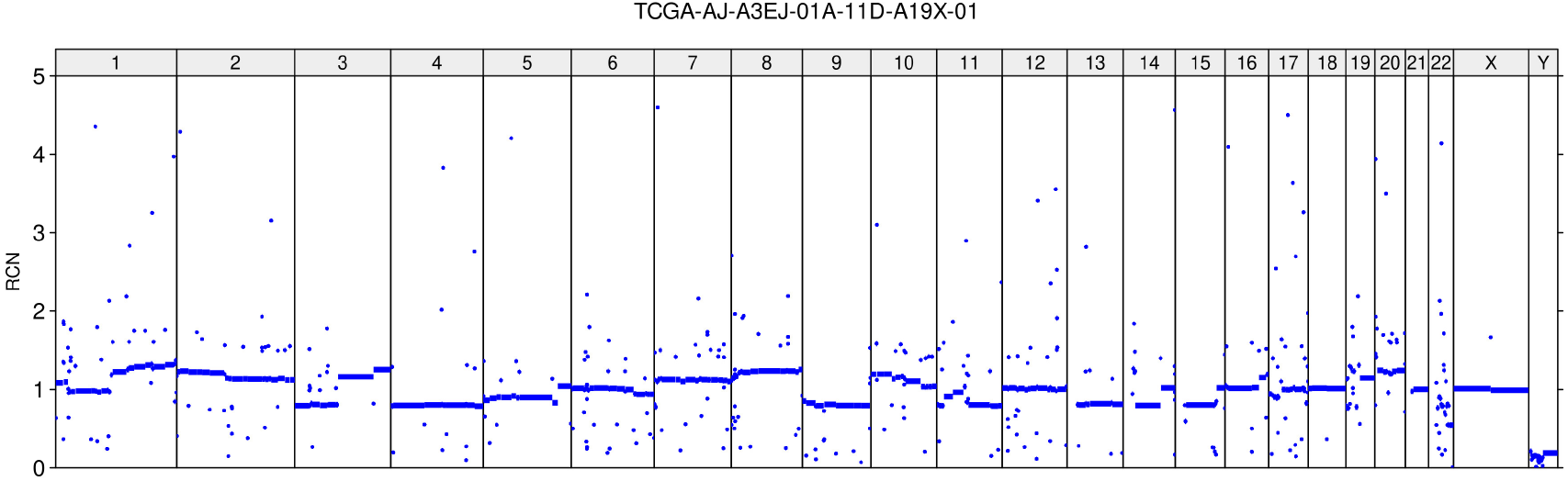
Relative copy number profile. Sample TCGA-AJ-A3EJ-01A-11D-A19X-01 from TCGA-UCEC study.

For this sample, the CNH value is 0.138437498026841, reported by the authors in the supplementary file 41467 2021 23384 MOESM4 ESM.xlsx, in row 9966 and column C, where `CNH_unfiltered` are stored. We obtained the same value of CNH by means of the matlab code provided by van Dijk et al. (2021), which can be accessed [here](https://github.com/dmmiedema/CNH). The code outputs also the estimated tumor purity 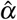, and the estimated tumor ploidy 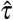. For this sample the values are 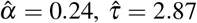. CNH estimates (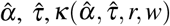 and 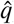) can also be obtained using the R library `rascal`; cf. Sauer et al. (2021).

For these values of 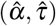 and the RCN profile of the sample, the resulting ACN profile 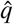, obtained using (*), is depicted on Fig. 2. Note the numerous segments with a negative number of copies.

**Figure 2.**
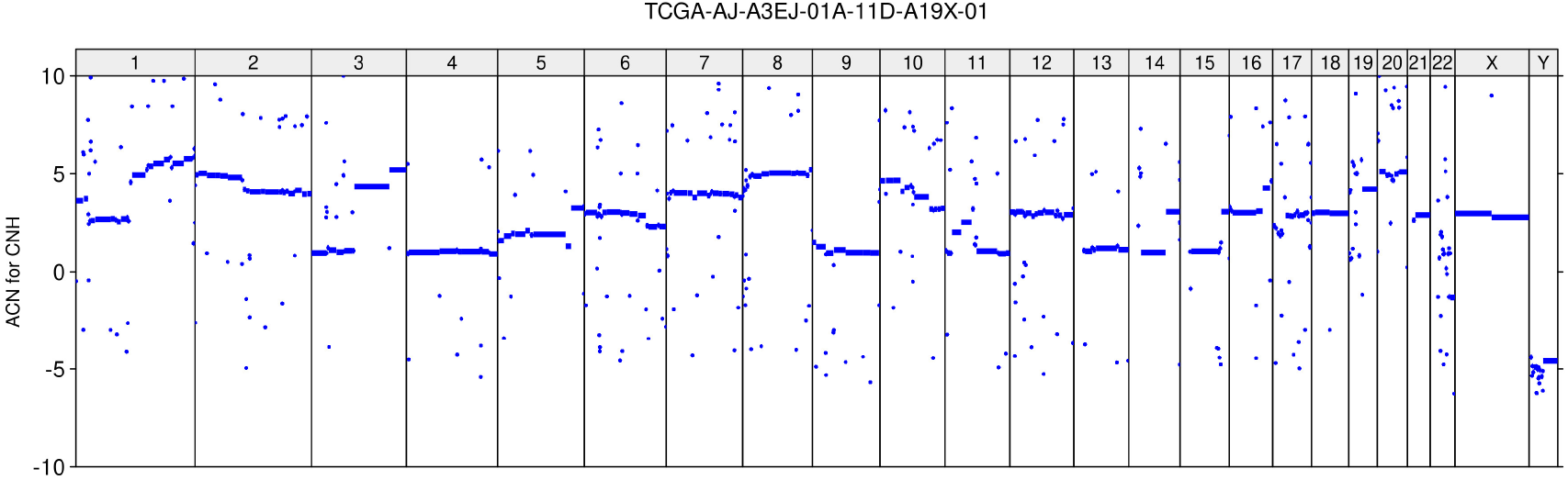
Absolute copy number profile recovered by CNH. Sample TCGA-AJ-A3EJ-01A-11D-A19X-01 from TCGA-UCEC study.

Could the case be saved by invoking the invariance of CNH? Let us see. The smallest value of 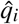 is −6.27 Hence, in order to make the shifted ACN profile 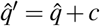 non-negative it would be necessary to shift it up by *c* = 6.27. The associated tumor ploidy 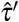 would be 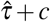, i.e., 9.14 which is well outside the range of values of tumor ploidy that van Dijk et al. (2021) consider as biologically meaningful in their search for CNH. Hence, for the sample TCGA-AJ-A3EJ-01A-11D-A19X-01 it is impossible to make the ACN profile of tumor which is non-negative and at the same time its CNH is 0.138437498026841.

For the sake of completeness let us look at the top 5 pairs of the tumor purity, tumor ploidy with the smallest values of *κ*; cf. Tab. (1). Note that the second-best pair is different than the best one and the remaining pairs are variation of the first two.

**Table 1.** Top 5 pairs of purity, ploidy with the associated value of *κ*; CNH method. Sample TCGA-AJ-A3EJ-01A-11D-A19X-01 from TCGA-UCEC study.

| purity | ploidy | kappa |
| --- | --- | --- |
| 0.24 | 2.87 | 0.1384375 |
| 0.27 | 3.86 | 0.1387208 |
| 0.24 | 2.86 | 0.1387284 |
| 0.27 | 3.87 | 0.1387558 |
| 0.24 | 2.88 | 0.1388188 |

The ANC profile for the 2-nd best pair (not shown) is ‘on average’ shifted by 1 up, relative to the 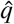 (exhibited on Fig. (1)) and consequently contains many segments with the negative number of copies.

As an aside, we note that the ACN profile which van Dijk et al. (2021) use to illustrate CNH (see Fig. 1g) in their Supplement (sample TCGA-AR-A24V-01A-21D-A166-01; file name EX-INE_p_TCGA_b142_SNP_N_GenomeWideSNP_6_B07_802436.hg19.seg.txt; [from GDC legacy](https://portal.gdc.cancer.gov/legacy-archive); CNH value 0.1035757; purity 0.32, ploidy 2.52) comprises 52 segments with the negative number of copies. They are not shown on the Fig. 1g) from the Supplement of van Dijk et al. (2021), since the y axis ranges from 0 to 8.

## 5 IMPOSING THE NON-NEGATIVITY OF ABSOLUTE COPY NUMBER PROFILE CONSTRAINT

The corrected definition of CNH, denoted CNHplus, - or CNH+, for short, - with the non-negativity constraint imposed on the absolute copy number profile, can formally be put as

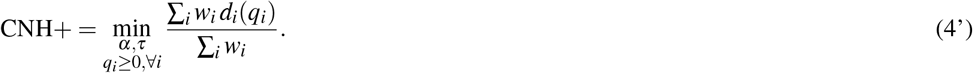

There, the notation *d_i_*(*q_i_*) is intended to stress that *d* depends on the ACN profile *q*. And *q*, in turn, depends on *α, τ* and *r*.

van Dijk et al. (2021) compute CNH over a grid, composed by the cartesian product of *α* = 0.2, 0.21,…, 1 and *τ* = 1.5, 1.55,…, 5. For a particular RCN profile *r*, for each pair (*α, τ*) of the grid the associated ACN profile can be computed by (*) and then fed to (·) to obtain the value of the objective function *κ*(*α, τ, r, w*). Afterwards, all (*α, τ*) for which the associated ACN profile contains at least one segment with negative copy number are excluded from the considerations. Sorting by value then leads CNH+.

For the TCGA-AJ-A3EJ-01A-11D-A19X-01 sample, CNH+is 0.1647427 and the estimated tumor purity is 1; the estimated tumor ploidy is 4.98. The resulting ACN profile, obtained using (*), is shown in Fig. 3.

**Figure 3.**
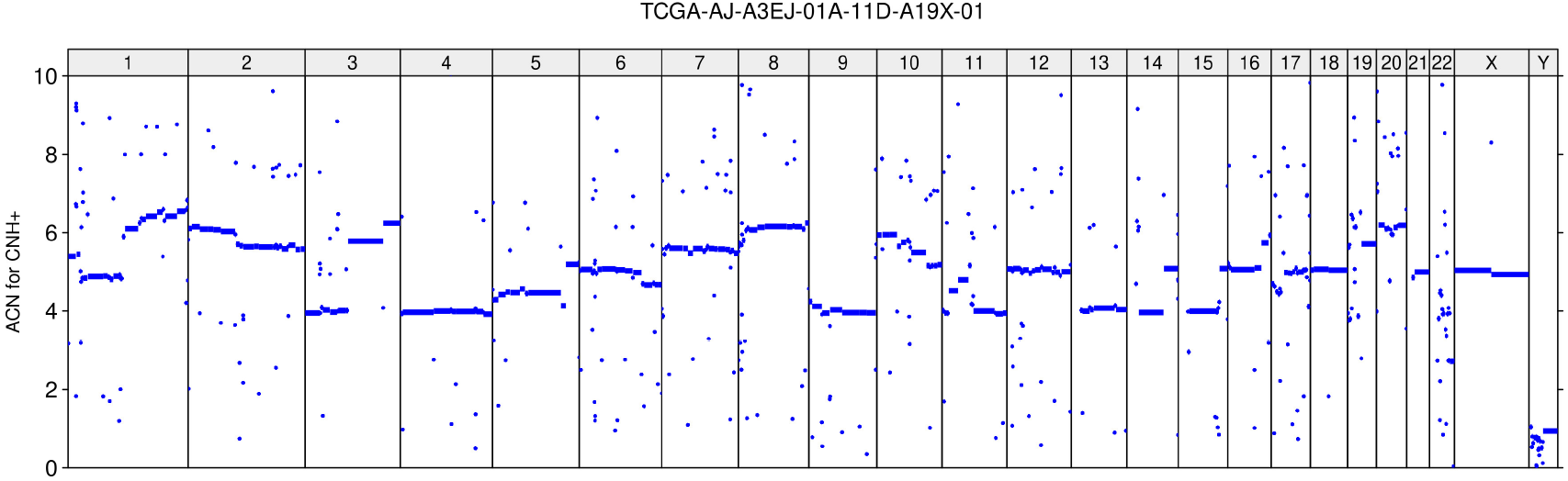
Absolute copy number profile recovered by CNH+. Sample TCGA-AJ-A3EJ-01A-11D-A19X-01 from TCGA-UCEC study.

The top 5 pairs of tumor purity, tumor ploidy ranked by CNH+can be found in the tab (2). The solution is unique.

**Table 2.** Top 5 pairs of purity, ploidy with the associated value of *κ*; CNH+method. Sample TCGA-AJ-A3EJ-01A-11D-A19X-01 from TCGA-UCEC study.

| purity | ploidy | kappa |
| --- | --- | --- |
| 1 | 4.98 | 0.1647427 |
| 1 | 4.97 | 0.1648025 |
| 1 | 4.96 | 0.1649735 |
| 1 | 4.95 | 0.1652086 |
| 1 | 4.99 | 0.1652292 |

## 6 OTHER SAMPLES FROM TCGA-UCEC STUDY

The ACN profile of the tumor obtained by CNH optimization was nonnegative for 106 out of the 540 tumor samples from the TCGA-UCEC study, which were considered by the authors. The remaining 434 samples have the CNH-recovered ACN profile such that at least one of its segments has a negative number of copies.

The cross-plot of the CNHplus values against the CNH values is shown in Fig. 4. The mean difference between CNHplus and CNH is 0.008; the maximal difference is 0.073. Of 540 TCGA-UCEC samples, 144 have CNHplus and CNH values apart by more than 0.01.

**Figure 4.**
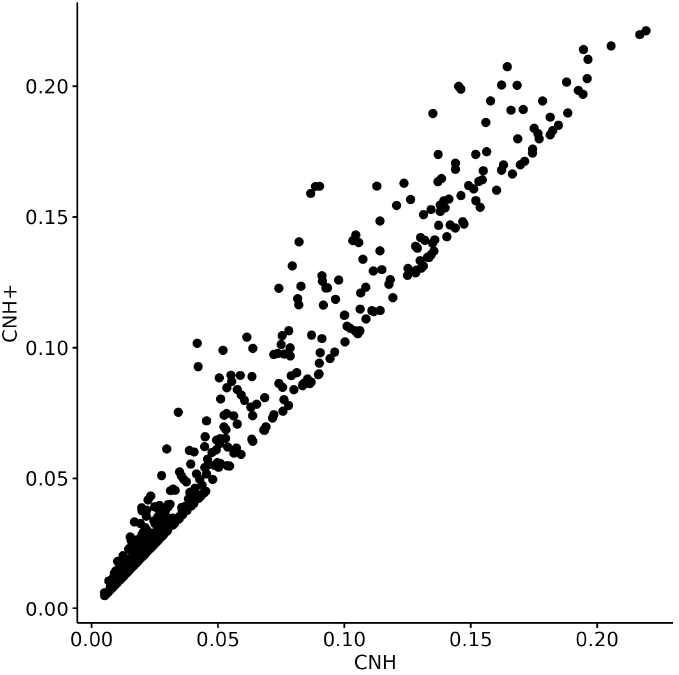
CNH+vs. CNH; TCGA-UCEC study.

CNH and associated 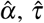 as well as CNH+ with associated values of tumor purity and ploidy are available at Supplement.

## 7 OTHER BIOLOGICAL CONSTRAINTS ON ABSOLUTE COPY NUMBER PROFILE

Besides non-negativity, the absolute copy number profile should satisfy other biological constraints as well.

First, the number of segments with zero copies cannot be excessively large, with the exception of chromosome Y. How this biological constraint should be implemented is an open issue. Possibly, the length of zero-copy segments or their number could be taken into account.

Another open problem concerns the sex chromosomes. Since their biology is quite different from that of autosomes, it would perhaps be reasonable to exclude them from consideration when computing CNH+.

Therefore, CNH+should not be considered as the final stage of correcting and improving the CNH measure. CNH+imposes the non-negativity of ACN profile, and corrects this flaw of CNH. The zero copies constraint and possibly other biological constraints could also be incorporated to CNH+, in order to provide a fully satisfactory CNH measure.

## 8 SURVIVAL STRATIFICATION BY CNH+

van Dijk et al. (2021) proposed CNH as a measure of ITH. For TCGA studies, the authors had split a patient cohort into two groups, by the median CNH value in the cohort. And they compared the two groups in terms of survival. The authors had found that for many types of cancer, the low CNH group and the high CNH group differ significantly in terms of survival.

The consequences of the deficiency in CNH for the findings of the authors can be assessed only indirectly. This is because the authors had used for the stratification of patients from TCGA studies the CNH based on RCN profiles with noise filtered out. In order to perform the noise filtering, it is necessary to have access to the probe-level copy number data, which is restricted; cf. Miedema (2022). This is why we have used above the unfiltered RCN profiles `CNH_unfiltered` to demonstrate the deficiency in CNH. And this is also why we can answer the question indirectly, only; using the unfiltered RCN profiles.

For the TCGA-UCEC study, Kaplan-Meier (KM) survival plots are exhibited on Fig. 5. For reference, the KM plot for `CNH` (i.e., CNH based on noise-filtered RCN profiles) is presented in panel A of the figure; reproducing Figure 4a from the paper. When comparing unfiltered CNH and unfiltered CNH+(Fig. 5, panels B, C), it can be seen that the effect of imposing the non-negativity of the ACN profile on survival is minor. Survival in the group of patients with CNH+ above its median value in the cohort is for the first 6 years better than for the respective group defined by `CNH_unfiltered`. The same can be said for the group below or equal to the respective median. The overall effect of using the corrected CNH on survival in TCGA-UCEC is minor. The conclusion of testing the null hypothesis of the equality of the two survival curves is not affected.

**Figure 5.**
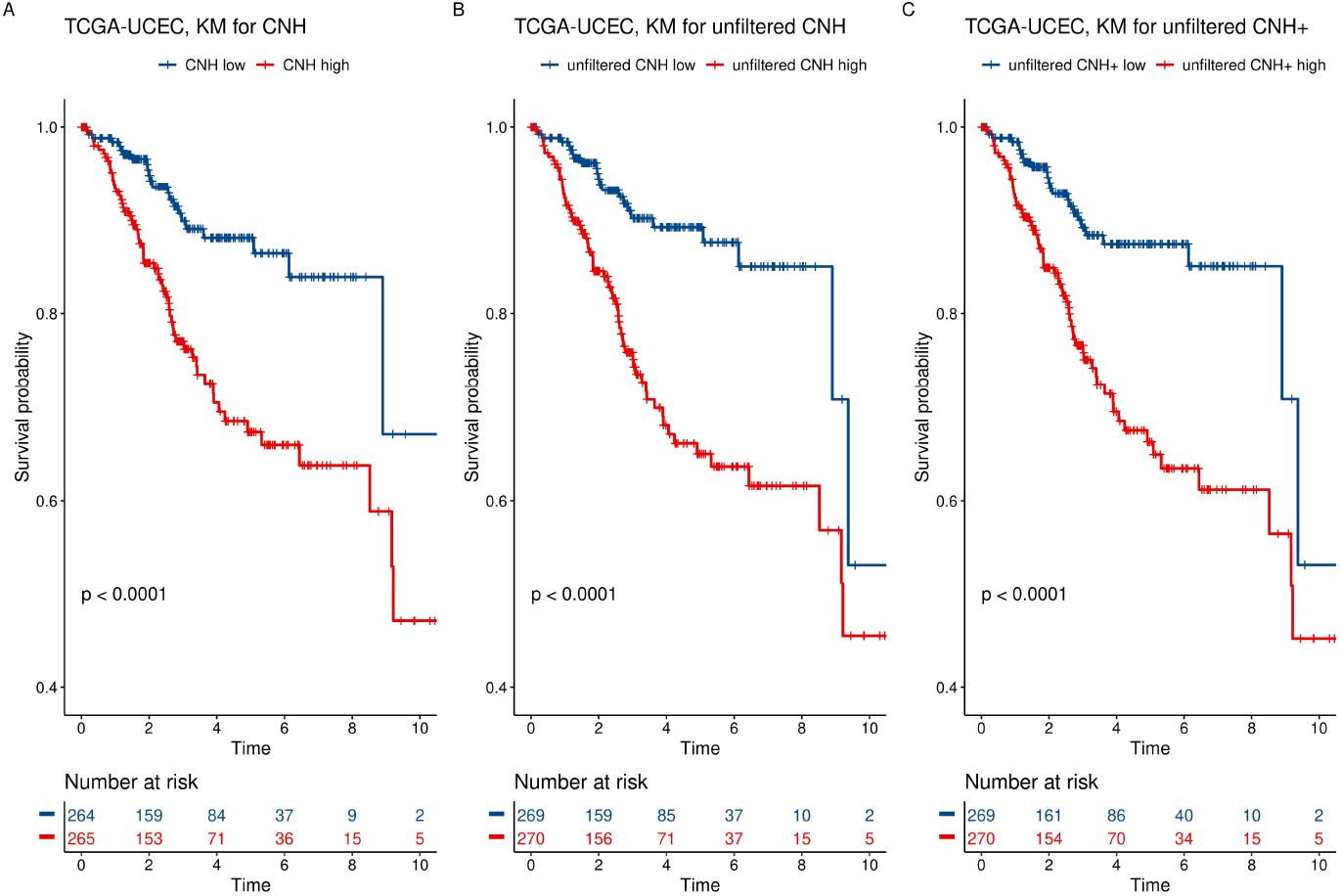
Kaplan-Meier fit to the overall survival data by group. A) CNH for noise-filtered relative copy number profiles, B) CNH for unfiltered relative copy number profiles, C) CNH+for unfiltered relative copy number profiles. TCGA-UCEC study.

TCGA-UCEC is one of the studies where CNH, as a tool for patient stratification, performed extremely well. For the TCGA testicular germ cell tumors (TGCT) study (see Shen et al. (2018)), the difference between the survival curves of the two cohorts defined by the median CNH (filtered) was not statistically significant; p-value was 0.31. (see Fig. 4 in the [Supplement](https://static-content.springer.com/esm/art%3A10.1038%2Fs41467-021-23384-6/MediaObjects/41467_2021_23384_MOESM1_ESM.pdf) to van Dijk et al. (2021) or Fig. 6A, here). Essentially the same stratification and the same p-value are obtained for CNH with the unfiltered RCN data; see Fig. 6B. However, the patient stratification based on CNH+(unfiltered) is radically different (see Fig. 6C) and the two cohorts split by median CNH+have statistically significantly different survival curves (p-value 0.044).

**Figure 6.**
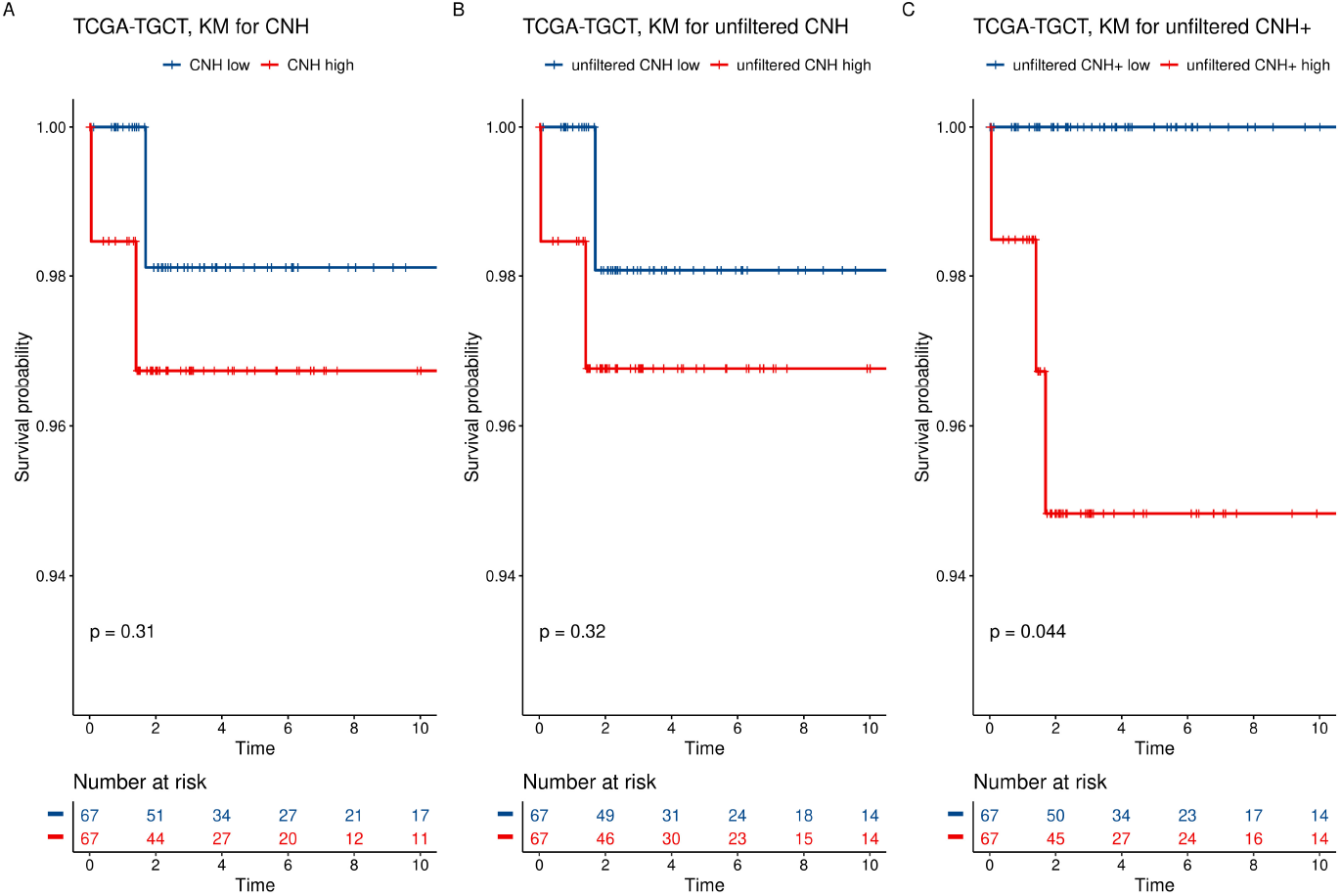
Kaplan-Meier fit to survival data by group. A) CNH for noise-filtered relative copy number profiles, B) CNH for unfiltered relative copy number profiles, C) CNH+ for unfiltered relative copy number profiles. TCGA-TGCT study.

For the TCGA Sarcoma Study (SARC), the survival stratification by CNH (filtered) leads to a p-value 0.015 and rather similar p-value (0.027) can be obtained for the stratification by CNH from the unfiltered RCN data. However, the stratification by CNH+(unfiltered) is not significant (p-value 0.14).

Kaplan-Meier survival curves (for CNH obtained from RCN profiles with noise excluded; for CNH obtained from unfiltered RCN profiles; for CNH+obtained from unfiltered RCN profiles) for all TCGA studies considered by van Dijk et al. (2021) can be found in Supplement.

## 9 CNH+VS. ABSOLUTE

ABSOLUTE (cf. Carter et al. (2012)) method for estimating the tumor purity, tumor ploidy, and recovering the ACN tumor profile from the RCN sample profile is usually considered as the reference method. ABSOLUTE is based on the same equations (2) and (1) that are used in CNH. However, Estimates are obtained in different way – ABSOLUTE uses an elaborate probabilistic model and its parameters are estimated by numeric optimization. ABSOLUTE assumes that cancer cells are dominated by a single clone, and unlike CNH, the method permits estimating also the tumor clonality.

Tumor purity and ploidy estimates by ABSOLUTE are available for the majority (89%) of the TCGA samples, considered by van Dijk et al. (2021). Therefore, it is of interest to compare CNH+estimates of tumor purity and ploidy with those of ABSOLUTE.

As can be seen in Fig. (7), ABSOLUTE tumor purity estimates differ dramatically from those obtained by CNH+. While ABSOLUTE estimates range from below 0.25 to 1, the CNH+estimates are predominantly close to 1, and do not reach much below 0.7. Although the Pearson correlation between ABSOLUTE and CNH+estimates of purity is statistically significant (p-value <0.0001), it is essentially zero (0.056). For CNH estimates of tumor purity, the correlation is 0.2 and the p-value <0.0001.

**Figure 7.**
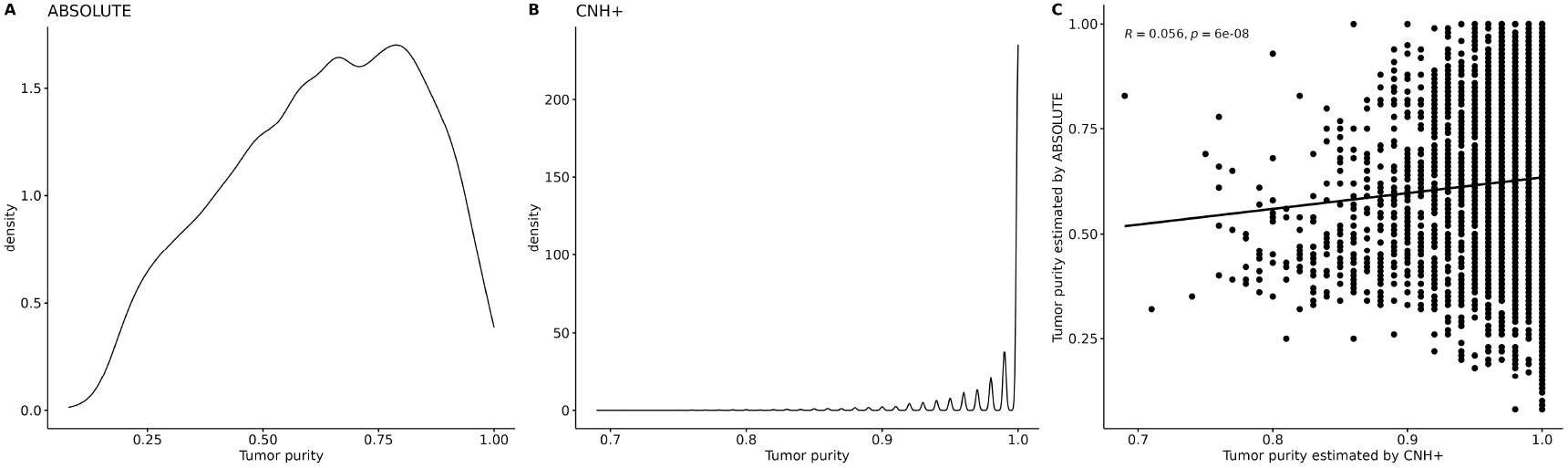
Estimates of the tumor purity by ABSOLUTE vs CNH+. A) Density plot of ABSOLUTE estimates of the tumor purity for TCGA samples. B) Density plot of CNH+estimates of the tumor purity for TCGA samples. C) Crossplot of ABSOLUTE estimates vs. CNH+estimates of the tumor purity for TCGA samples. Pearson correlation coefficient *R* and the p-value for the test of the null hypothesis that the population correlation is zero are at the upper-left corner. The ABSOLUTE estimates of tumor purity were taken from the Pan-Cancer Atlas, file TCGA-CDR-SupplementalTableS1.xlsx available at https://gdc.cancer.gov/about-data/publications/pancanatlas

Not surprisingly, ABSOLUTE estimates of the average tumor ploidy are of different nature than those of CNH+; see Fig. (8). ABSOLUTE estimates of tumor ploidy are predominantly close to 2 and other values are rather rare. CNH+estimates of tumor ploidy equal to 3 are only around three times less frequent than those equal to 2.

**Figure 8.**
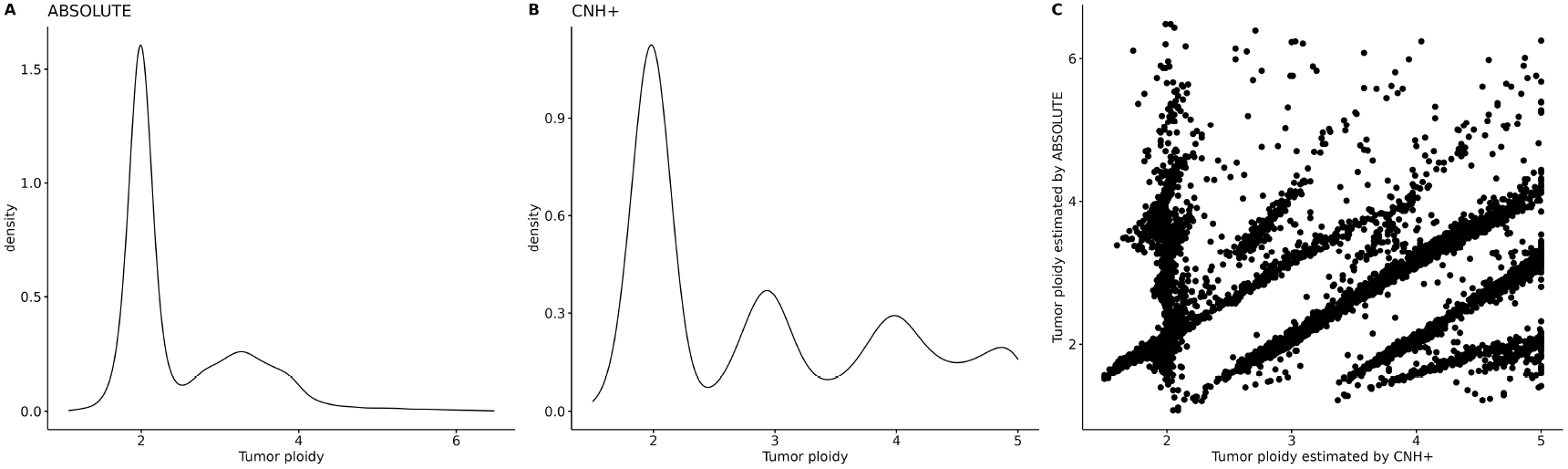
Estimates of the tumor ploidy by ABSOLUTE vs CNH+. A) Density plot of ABSOLUTE estimates of the tumor ploidy for TCGA samples. B) Density plot of CNH+estimates of the tumor ploidy for TCGA samples. C) Crossplot of ABSOLUTE estimates vs. CNH+estimates of the tumor ploidy for TCGA samples. The ABSOLUTE estimates of tumor ploidy were taken from the Pan-Cancer Atlas, file TCGA-CDR-SupplementalTableS1.xlsx; https://gdc.cancer.gov/about-data/publications/pancanatlas

Recent literature suggests that ABSOLUTE substantially underestimates tumor purity. Poell et al. (2019) compared ABSOLUTE estimates of tumor purity with the gold standard in 253 ovarian cancer samples; the result can be found here. Sauer et al. (2021), see their Fig. 1, used ovarian cancer cell lines and patient-derived xenograft with known high purity to evaluate purity estimation by ABSOLUTE and two other methods (ichorCNA and ACE). As it can be seen from Fig. 1b) in Sauer et al. (2021), ABSOLUTE substantially underestimates the tumor purity; as do also the other two methods, but to a lesser extent.

Haider et al. (2020) used prostate cancer samples from the TCGA-PRAD study to compare tumor purity estimates obtained by a ten bioinformatic methods with estimates made by pathologists. The authors concluded that there is a poor concordance between pathological and bioinformatic purity estimates. Interestingly enough, the tumor purity estimates obtained by LEUC, one of the methods considered by the authors, which uses the genomic methylation signature of leukocytes, were concentrated around 0.9, similar to CNH+estimates.

## 10 DISCUSSION

CNH, introduced by van Dijk et al. (2021), is an interesting single-sample method for quantifying ITH from copy number variation data. If the RCN profile of the tumor sample is available, then the ACN profile of the tumor is recovered by CNH, together with the associated estimates of the tumor purity and the average tumor ploidy. The estimates are obtained from a minimization problem, where the objective function is the weighted distance of the tumor ACN profile to integer values. CNH is just the value of the objective function at the minimum. The authors used CNH to stratify cancer patients for survival.

The recovered ACN profile of the tumor should, in our view, satisfy biological constraints. Unfortunately, restricting tumor purity and tumor ploidy in the CNH optimization to biologically meaningful values does not, in general, guarantee that the recovered ACN profile of tumor is nonnegative in every segment. There may be segments with the negative number of copies, as illustrated in the TCGA-UCEC study above. CNHplus (CNH+, for short), the corrected CNH method, which respects the non-negativity of the tumor ACN profile, was introduced here as a step towards obtaining the ITH measure which satisfies key biological constraints. This is why it can also be used to develop numerical characteristics other than copy-number heterogeneity, such as copy number signatures (cf. Steele et al. (2022)) or various Copy Number Alteration scores (cf. Franch-Expósito et al. (2020)), which can be added to predictive modeling of patient survival.

The effect of using CNH versus CNHplus on the stratification of cancer patients for survival can be minimal (as shown for endometrial cancer - UCEC study) or significant (for testicular germ cell tumors - TGCT study, or sarcomas – SARC study). Further, restricting the number of segments with zero copies appears, in our view, to be the next constraint to consider in future developments of CNHplus.

## 11 ACKNOWLEDGEMENTS

MG sincerely thanks Daniël M. Miedema and Erik van Dijk for their patient answering numerous questions about details of CNH. The results shown here are in whole based upon data generated by the TCGA Research Network: https://www.cancer.gov/tcga.

